# Negative curvature-promoting lipids instruct nuclear ingression of low autophagic potential vacuoles

**DOI:** 10.1101/2021.06.26.450031

**Authors:** Manon Garcia, Sylvain Kumanski, Alberto Elías-Villalobos, Caroline Soulet, María Moriel-Carretero

**Affiliations:** Centre de Recherche en Biologie cellulaire de Montpellier (CRBM), Université de Montpellier, Centre National de la Recherche Scientifique, Montpellier, France; Institut Curie, UMR3244, Paris, France

**Author notes:** equal contribution (Manon Garcia, Sylvain Kumanski).

## Abstract

Membrane contact sites are functional nodes at which organelles exchange information through moving ions, proteins and lipids, thus driving the reorganization of metabolic pathways and the adaptation to changing cues. The nuclear-vacuole junction of *Saccharomyces cerevisiae* is among the most extensive and better-known organelle contact sites, described to expand in response to various metabolic stresses. While using genotoxins with unrelated purposes, we serendipitously discovered a phenomenon that we describe as the most extreme and intimate contact ever reported between nuclei and vacuoles: the vacuole becomes completely internalized in the nucleus. We define lipids supporting negative curvature, such as phosphatidic acid and sterols, as *bona-fide* drivers of this event. Functionally, we purport that internalized vacuoles are low efficiency ones whose removal from the cytoplasm optimizes cargo interaction with functional vacuoles. Thus, our findings also point to nucleus-vacuole interactions as important for metabolic adaptation. Yet, rather than by inter-organelle exchanges, the underlying mechanism literally concurs with vacuolar sequestration.

## Introduction

Eukaryotic cells possess a functionally committed system of endomembranes whose regulated remodeling is essential to warrant adaptation to stresses, changing cues and cell cycle requirements. Among them, the Endoplasmic Reticulum (ER) is one of the most dynamic, suffering drastic transitions during ER stress, when the volume of membranes massively expands to increase its protein folding capacity (1). The perinuclear subdomain of the ER, also known as the nuclear membrane, is particularly prone to extreme remodeling. Irrespective of nuclear division occurring in an “open” or in a “closed” manner, the perinuclear ER membranes will undergo dramatic changes either because of rupture, dispersion and re-assembly, or because of expansion and deformation (2,3). An important aspect of membrane remodeling concerns the sites of physical proximity between different endomembrane systems, known as membrane contact sites (4). At these locations, membranes belonging to two different organelles, such as the ER and the Golgi, or the mitochondria and the Lipid Droplets, stay in close proximity (10 to 80 nm), which allows the spatial organization of enzymes involved in a given metabolic pathway (5), as well as the active exchange of different molecules, such as lipids and ions (6).

A membrane contact site that can become impressively extensive is that between the nucleus and the vacuole (the equivalent to the lysosome) of the yeast *Saccharomyces cerevisiae*, termed the Nucleus-Vacuole Junction (NVJ). The tether between these two membranes is actively maintained by the proteins Vac8 and Nvj1, present in the vacuolar and outer nuclear membranes, respectively (7). Helped by additional factors, such as Snd3 (8), the NVJ expands “zipper-wise” during nutritional stress, such as glucose and aminoacid shortage, and upon Target Of Rapamycin Complex (TORC) inhibition (9). This increase in the contact surface can serve to send esterified lipids for storage within Lipid Droplets (9), to increase the flux of metabolites through the mevalonate pathway (5) or to recycle non-essential nuclear components (10). This latter process transfers components mostly arising from the nucleolus directly to the vacuole without the need of transporting vesicles (the autophagosomes). This way, when the ribosomal DNA in the nucleolus stops being actively transcribed as to spare energy, the vacuole will degrade both nucleolar proteins that promote active transcription as well as forming ribosomes, thus helping match a decrease in translation capacity (10–12).

But the NVJ is not only important to respond to nutritional shortage. Recently, it has been shown that the consequences derived from compromising the assembly of nuclear pore complexes at the nuclear membrane are alleviated by increasing the membrane contact sites between the nucleus and the vacuole (13). Moreover, the NVJ is a site for the synthesis of long chain fatty acids that impacts the sphingolipid biosynthetic pathway (14). As such, lack of appropriate membrane tethering at this location sensitizes cells to sphingolipid synthesis inhibitors even under basal conditions (15). Since it is emergently recognized that membrane contacts sites are pro-active in exchanging lipids and signals between organelles (6), it stems that the contact between the nucleus and the vacuole may still have multiple secrets to deliver.

In this work, we define lipid scenarios in which the nucleus and the vacuole interact in the most extreme manner described up-to-here in the literature. We find that enrichment at membranes of phosphatidic acid or free sterols promote the internalization of the vacuole within the nucleus. We note that these lipids support negative curvature of membranes, which may be key during the invagination process. Yet, given their fusogenic potential, the phenomenon could be alternatively taking place through membranes fusion. Whichever the case, we asked whether this concomitant loss of vacuoles from the cytoplasm impacted general (macro)autophagy of cytoplasmic cargoes and found, to our surprise, that autophagy is favored this way. We provide arguments to propose that internalized vacuoles are low efficiency ones whose removal from the cytoplasm optimizes cargo interaction with functional vacuoles. Our results therefore unveil an unprecedented membrane-remodeling event with direct impact in metabolic adaptation. Further, our findings question how the invasion of the nucleoplasmic space by such voluminous bodies affects genome homeostasis.

## Results

### 1. Identification of unusual structures inside the nucleus during treatment with different genotoxic agents

In order to undoubtfully identify the nucleus when monitoring the formation of foci by DNA repair factors, we routinely use the nucleoplasmic protein Pus1 tagged with mCherry in its N-terminal region (Figure S1A). We realized that, when some genotoxic agents are added to cell cultures growing exponentially in rich medium, the Pus1 signal in some nuclei became displaced, or even absent, giving rise to what we informally defined as “holes” (Figure 1A, and Figure S1B). This serendipitous but repetitive and striking observation prompted us to investigate the nature of such structures. We therefore systematically quantified the percentage of cells displaying at least one of these holes upon exposure to different genotoxic agents. We found that, already basally, 6% of the cells manifested this phenomenon. Methylmethanosulfonate (MMS) modestly doubled this percentage and, most pronouncedly, zeocin triggered a time-dependent increase in the frequency of these structures (Figure 1B). On the contrary, 4-Nitroquinoline 1-oxide (4-NQO), camptothecin (CPT) and hydroxyurea (HU), genotoxins that affect DNA differently, did not induce the formation of these structures (Figure 1B). All the genotoxic agents used in this set-up provoke cells arrest in different stages of the cell cycle, which validated their activity (Figure 1C, red asterisks). These data suggest that the “nuclear holes” phenotype could be unrelated to DNA damage. Zeocin has been reported to provoke an arrest in G2 during which DNA segregation towards the daughter cell is paused while nuclear membrane expansion continues, leading to the accumulation of overgrown nuclear membranes (23). This phenotype is not triggered by HU, as in our case, but it is strongly elicited by nocodazole. We therefore tested whether nocodazole, a microtubule-depolymerizing drug, also induces nuclear holes formation, and we found it did robustly (Figure 1D). Both nocodazole and zeocin lead to the accumulation of cells in G2/M phases of the cell cycle and, as estimated from the cytometry profiles, part of the MMS-treated cells also reaches this cell cycle stage during our experimental framework (Figure 1C), suggesting that this could be a necessary trigger. Yet, it is manifestly not sufficient, for cells treated with CPT or 4-NQO also accumulate in G2/M (Figure 1C) without displaying “nuclear holes” (Figure 1B, S1B). Thus, the formation of nuclear holes is elicited preferentially (but not sufficiently) in G2/M and does not seem to require DNA damage *per se*.

**Figure 1.**
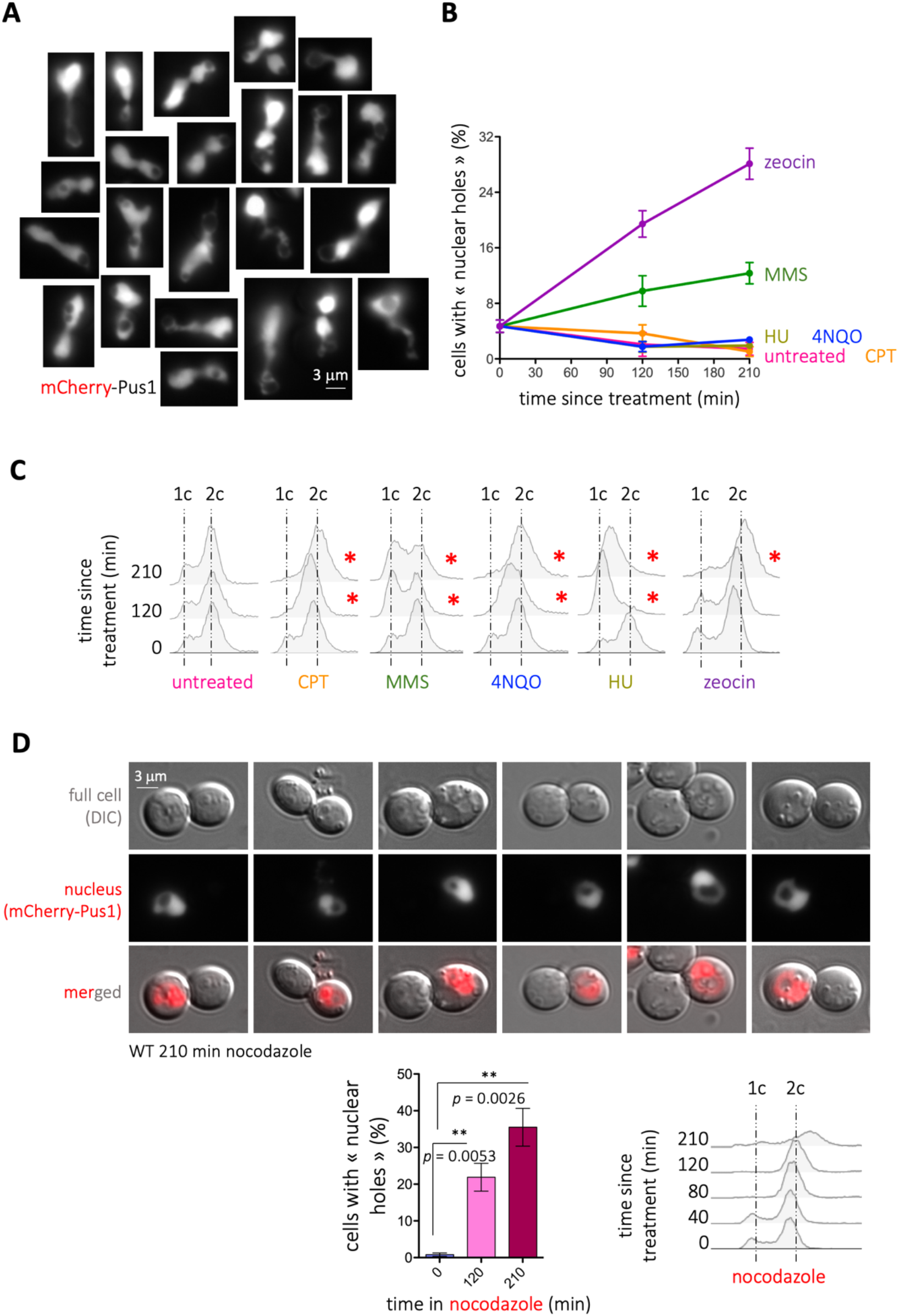
Detection of nuclear holes in response to genotoxic agents. **(A)** Illustrative images of mCherry-Pus1 signals (nuclei) from *Saccharomyces cerevisiae* cells exposed to 100 μg/mL zeocin for 210 min in which “black holes” can be observed. Eventual saturated images are so to permit the delineation of the Pus1 signal surrounding the holes. **(B)** Quantification of the percentage of cells displaying nuclear holes in response to the indicated genotoxic agents at the indicated time-points. The used doses were 100 μg/mL zeocin, 100 mM HU, 0.1% MMS, 100 μM CPT and 0.05 mg/L 4-NQO. The plotted values and the error bars are the mean and the Standard Error of the Mean (SEM), respectively, of 3 independent experiments. At least 200 cells were counted per time-point, treatment and experiment. **(C)** Cytometry profiles of the experiment shown in (B). 1c and 2c indicate the DNA content. Red asterisks mark the time-points when alterations in the cell cycle profiles can be detected, as compared to the untreated samples. **(D)** WT cells bearing the mCherry-Pus1 construct were grown in rich medium to the exponential phase and exposed to 15 μg/mL nocodazole for the indicated time. Cells were imaged and 6 examples are shown. The percentage of cells displaying nuclear holes was calculated and is plotted. The bars and the error bars are the mean and the SEM, respectively, of 3 independent experiments. At least 200 cells were counted per time-point, treatment and experiment. The *p*-values indicate the statistical significance upon performing a t-test. Cytometry profiles are shown, where 1c and 2c indicate the DNA content

### 2. Phosphatidic acid supports the formation of zeocin-triggered nuclear holes

Given the strength of the phenotype, we focused on the treatment with zeocin to further characterize the phenomenon of nuclear holes formation. As cell cycle progression relates to nutrient availability, we first tested whether nutritional conditions could have any impact on the apparition of these structures. Importantly, in marked contrast to what we observed in rich medium, growing the cells in defined, minimal medium abolished the formation of nuclear holes in response to zeocin (Figure 2A, pEmpty). Of note, zeocin was efficiently incorporated in minimal medium-growing cells, as demonstrated by the ability of the DNA damage sensing factor Tel1 to form zeocin-induced foci (Figure S1C). Together, this suggests that a factor(s) needed to form the holes, presumably abundant in rich medium, may become limiting in minimal medium. Although the origin of the holes could be diverse, it was likely that they derive from transitions at the nuclear membrane. Among the many molecules needed to remodel membranes whose availability is impacted by nutritional conditions are lipids. In that context, lipids supporting negative curvature (Figure 2B, yellow phospholipids), such as Phosphatidic Acid (PA) or DiAcylGlycerol (DAG) (24,25), are expected to be relevant. Overexpression for 2 h of the PA-generating enzyme Dgk1 (Figure 2C) led to an imperceptible increase in the presence of nuclear holes (Figure 2A, p*DGK1*^OE^, time “0”). Yet, maintaining the overexpression 210 min longer led to 35% of cells displaying nuclear holes (Figure 2A, p*DGK1*^OE^, time “210*”). Moreover, addition of zeocin to Dgk1-overexpressing cells doubled the percentage of cells displaying nuclear holes in only 20 min, despite growth occurring in minimal medium, and led to a final 55% of cells bearing the phenotype (Figure 2A, p*DGK1*^OE^, times “20-210”). On the contrary, overexpression of the hyper-active Pah1 allele *Pah1-7A*, which promotes the accumulation of DAG at the expense of PA (Figure 2C), did not provoke any increase in the percentage of cells displaying nuclear holes, even after 210 min since zeocin addition (Figure 2A, p*PAH1-7A*^OE^). These data point at PA as relevant to form nuclear holes, and exclude DAG molecules as implicated in this process, in spite of their conical shape promoting negative curvature (24). In support, the reciprocal approach using deletion mutants demonstrated that an excess of PA creates a constitutive presence of nuclear holes in approximately half of the population (Figure 2D, *pah1Δ*), but the chronic excess of DAG only slightly yet not significantly differed from the isogenic WT strain (Figure 2D, *dgk1Δ*).

**Figure 2.**
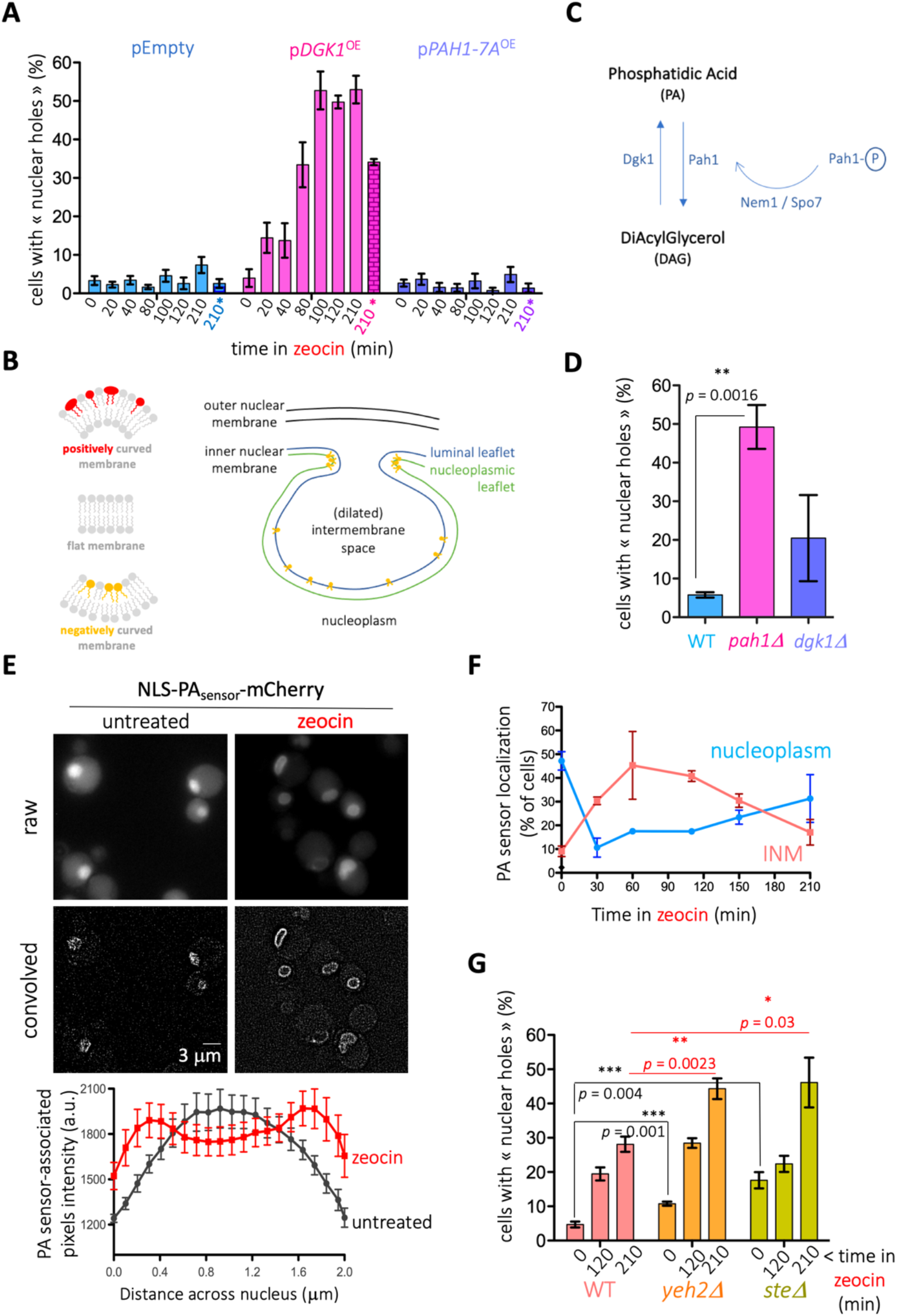
Lipid determinants of nuclear holes formation. **(A)** Cells bearing the genomic mCherry-Pus1 construct were grown overnight in minimal medium selective for the indicated plasmids with glycerol as the carbon source. The exponential cultures were then supplemented with 2% galactose to induce the expression of nothing (pEmpty), of Dgk1 (p*DGK1*^OE^) or of the hyperactive Pah1-7A (p*PAH1-7A*^OE^). Two hours later (time “0”), 100 μg/mL zeocin was added. The indicated time-points therefore indicate the elapsed time since zeocin addition, with the exception of the last point of each set (“210*” and marked in color), which accounts for the impact of the overexpression only. The percentage of cells in the population displaying at least one nuclear hole was counted. Each bar reflects the mean of 3 independent experiments, and the error bars account for the SEM. At least 200 cells were considered per time point, condition and experiment. **(B)** Left: simplified scheme of basic membrane curvature set-ups in which conical phospholipids (in yellow) help shape membranes of negative curvature, cylindrical phospholipids (in grey) give rise to flat membranes and inverted conical phospholipids (in red) serve to shape positively curved membranes. Right: invaginations (for simplicity only of the inner nuclear membrane, INM) request negative curvature-promoting phospholipids (in yellow). This requirement is modest at regions such as the luminal leaflet of the INM, while it is maximal at nascent sites in the nucleoplasmic leaflet. **(C)** Simplified scheme illustrating the enzymes responsible for PA and DAG synthesis. Pah1 is subjected to inactivation by phosphorylation (Pah1-P). To bypass the need of the phosphatase Nem1/Spo7 complex in order to activate it, we have overexpressed a constitutively de-phosphorylated isoform, *Pah1-7A*. **(D)** Cells of the indicated genotypes bearing the genomic mCherry-Pus1 construct and growing exponentially in rich medium were photographed and the percentage of cells in the population displaying at least one nuclear hole was counted. Each bar reflects the mean of 3 independent experiments, and the error bars account for the SEM. At least 200 cells were considered per condition and experiment. The *p*-value indicates the statistical significance upon performing a t-test. **(E)** (Top) Exponentially growing (in rich medium) WT *S. cerevisiae* cells transformed with a previously validated, nucleus-directed, mCherry-tagged sensor capable of detecting membrane-bound Phosphatidic Acid (NLS-PA_sensor_-mCherry (21), derived from the Q2 domain of Opi1 (56)) were treated (or not) with 100 μg/mL zeocin and inspected by fluorescence microscopy. Representative raw images or its processed counterparts after using the “convolve” tool in ImageJ are displayed. “zeocin” image belongs to timepoint 110 minutes of (F). Please note that different intensities among cells may be due to the biosensor being expressed from a plasmid. (Bottom): a line was drawn through nuclei using the raw images, and pixel intensity values across the line (distance) were plotted for both zeocin (red line) and untreated (black line) conditions. The graph displays the mean intensity and the SEM for n = 7 at timepoint 110 minutes. **(F)** The same WT cells illustrated in (E) were followed in time after zeocin addition. The percentages of cells displaying either nucleoplasmic (blue line) or perinuclear (INM: inner nuclear membrane, pink line) localization of the PA-associated signal are plotted. Please note that the addition of both nucleoplasmic and perinuclear percentages does not reach 100 %. This is due to the presence in the population of cells displaying either lack of signal, or vacuolar signal (presumably due to sensor degradation). The plotted values are the mean and the SEM of 3 independent experiments. **(G)** Cells of the indicated genotypes bearing the genomic mCherry-Pus1 construct and growing exponentially in rich medium were photographed before 100 μg/mL zeocin addition, and 120 and 210 min later. The percentage of cells in the population displaying at least one nuclear hole was counted. Each bar reflects the mean of 4 to 7 independent experiments, and the error bars account for the SEM. At least 200 cells were considered per condition and experiment. The *p*-values indicate the statistical significance upon performing the annotated t-tests.

If PA is important to trigger transitions at the nuclear membrane in response to zeocin, then zeocin treatment is expected to trigger PA accumulation or redistribution, at least at the inner nuclear membrane (INM). We used a nucleus-targeted fluorescent PA biosensor (description in the legend of Figure 2E), whose correct localization could be determined by its basal nucleoplasmic diffuse signal (Figure 2E, top left). We then treated cells growing exponentially in rich medium with zeocin and monitored the localization of the PA biosensor signal in time. Importantly, the biosensor signals became perinuclear (Figure 2E, top right), peaking at 60 to 100 min after zeocin treatment (Figure 2F). Later on, the PA biosensor progressively became nucleoplasmic again, indicative of PA detection at the INM being transient (Figure 2F). As a control for the specificity of this behavior, we were unable to observe this transient signal enrichment at the INM if the experiment was performed in minimal medium. Thus, PA seems to be a key molecule in promoting the formation of nuclear holes in general, and in response to zeocin in particular.

### 3. Sterols co-operate in the formation and maintenance of nuclear holes

Apart from PA or DAG, sterols have demonstrated to promote negative curvature at biological membranes (25). In order to increase the concentration of sterols in membranes and assess their putative contribution to nuclear holes formation, we used mutant strains either impaired in the storage within cytoplasmic LD of free sterols (*are1Δ are2Δ,* simplified in the literature as *steΔ*) or unable to release sterols from membranes (*yeh2Δ*) (26). Importantly, both mutants modestly but reproducibly displayed nuclear holes in a basal manner (Figure 2G). Addition of zeocin led to an additive formation of nuclear holes (Figure 2G). Thus, membrane-embedded sterols are not needed to form nuclear holes in response to zeocin yet can act as adjuvants, perhaps by favoring a membrane context that is more susceptible to their genesis.

### 4. Nuclear holes do not necessarily associate to the nucleolus

The structures under study could therefore relate to invaginations from the inner or from the whole nuclear membrane, or even fusion events with cytoplasmic structures. Since the part of the nuclear membrane associated to the nucleolus is repeatedly reported as prone to expansion and to support lipid transitions (23,27–29), we next assessed the position of nuclear black holes with respect to the nucleolus. To do so, we used two fluorescent nucleolar markers: either Nop1-CFP, a protein soluble in the nucleolus (19); or Net1-GFP, a ribosomal DNA-bound factor (20). We found that, irrespective of the marker, the nucleolus was frequently close to the holes (Figure 3A). Yet, the percentage of cells displaying the black hole inside the nucleolus (i.e. the hole was irrefutably in the middle of the nucleolar signal) accounted for a small percentage of all the events, and this even in genetic contexts where black holes were very frequent, such as in the *pah1Δ* strain, or in its genetic mimic *nem1Δ* (Figure 3B). Thus, either the formation of the holes does not necessarily relate to the nuclear membrane adjacent to the nucleolus, or it does but holes display mobility that makes them diffuse away from this nuclear subdomain.

**Figure 3.**
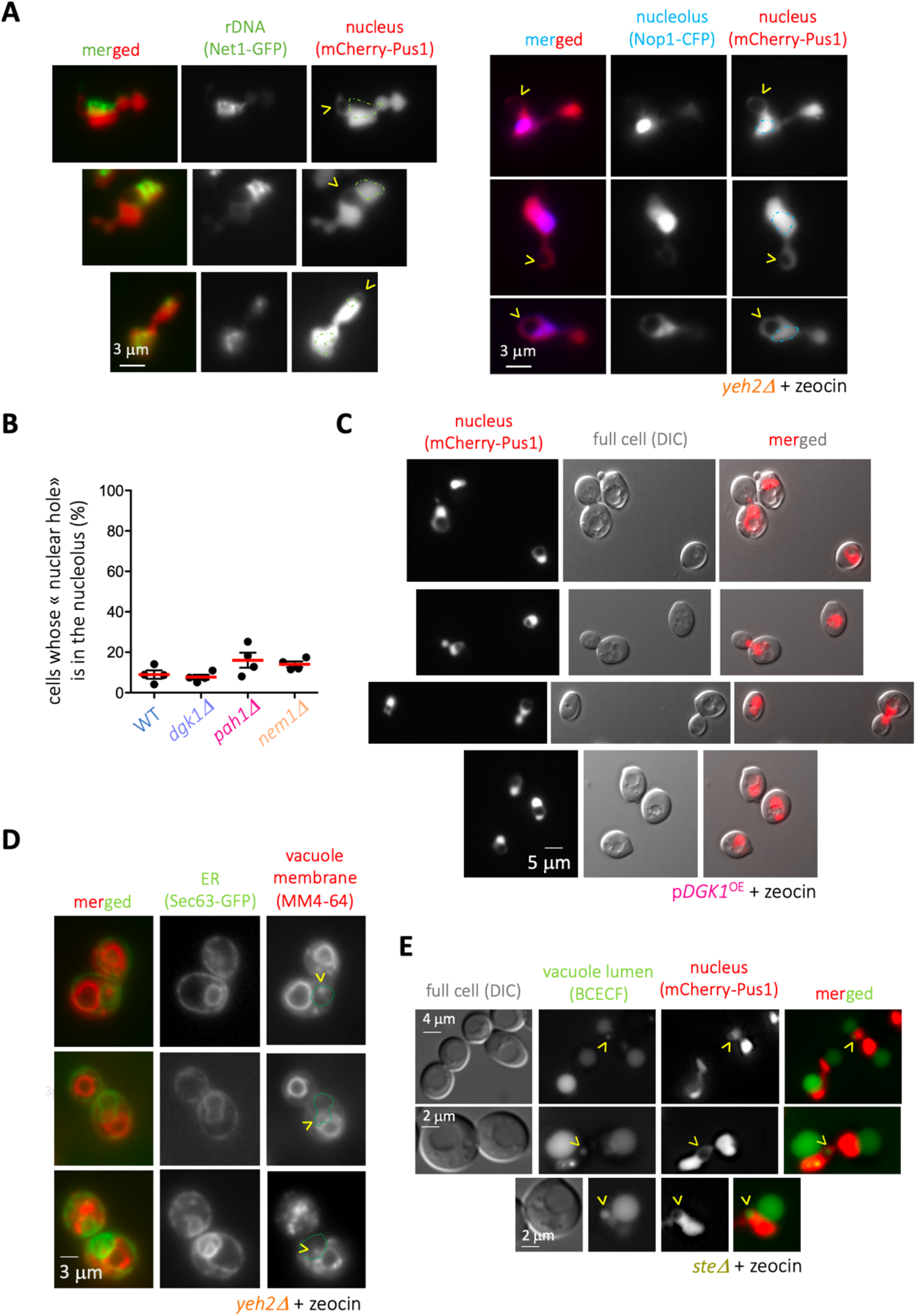
Determination by microscopy of the nature of the nuclear holes. **(A)** The strain *yeh2Δ*, which displays nuclear holes at high frequency in response to 100 μg/mL zeocin (Figure 2G), was transformed either with a vector expressing Net1-GFP, to mark the position of the ribosomal DNA, or with a vector expressing Nop1-CFP, to mark the position of the nucleolus, grown to the exponential phase and treated with that drug. The comparison with nucleoplasmic mCherry-Pus1 signals allows to monitor the relative position of the nuclear holes with respect to these sub-nuclear domains. To facilitate visualization, the contour of the rDNA or the nucleolar signals has been over-imposed onto the nucleoplasmic ones. **(B)** The indicated strains, transformed with the vector expressing Nop1-CFP, were monitored for the relative position of the nuclear holes with respect to the nucleolus. The percentage of cells in the population in which the nuclear hole disrupted the nucleolus was counted. The red bar reflects the mean of 4 independent experiments, and the error bars account for the SEM. At least 200 cells were considered per condition and experiment. **(C)** Selected examples of cells in which the comparison of the mCherry-Pus1 signals and the position of the vacuoles, as inferred from the differential interference contrast (DIC) images, permit to infer that the nuclear holes correspond to the vacuole. **(D)** The strain *yeh2Δ*, which displays nuclear holes at high frequency in response to 100 μg/mL zeocin (Figure 2G), was transformed with a vector expressing Sec63-GFP to mark the Endoplasmic Reticulum, which includes the nuclear membrane. Exponentially growing cells treated with this drug for 210 min were dyed with the vacuole membrane marker MM4-64 and imaged. The rationale was to try to identify vacuoles inside the nucleus. Apart from doubtful events due to the focal plane, these events were rare. Three examples in which vacuole-reminiscent bodies, dyed with MM4-64, could be found inside the nucleus, are shown. To facilitate comparison, the nuclear contour has been over-imposed onto the MM4-64 channel and pointed at by yellow arrowheads. **(E)** The strain *steΔ*, which displays mCherry-Pus1-defined nuclear holes at high frequency in response to 100 μg/mL zeocin (Figure 2G), was grown in rich medium to the exponential phase and treated with this drug for 3 hours. Vacuole lumens were dyed using the dye BCECF and cells immediately imaged. Yellow arrowheads point at nuclear holes dyed with the BCECF marker.

### 5. Nuclear holes correspond to nucleus-engulfed vacuoles

We next aimed at understanding the nature of the nuclear holes. A re-inspection of the images made it apparent that the black holes corresponded to vacuoles in the DIC channel (Figure 3C). Importantly, these examples neatly differed from situations where the vacuole is so big that it pushes, therefore deforms, the nucleus (Figure S2A). By using the vacuole membrane-specific dye MM4-64, which emits in the red wavelength range, we could validate that the nuclear holes adjacent to or within the Nop1-CFP signals were vacuoles (Figure S2B). However, some nuclear black holes that could be inferred adjacent to the Nop1 signal were partially refractory to MM4-64 staining (Figure S2B, arrowheads). These data suggested that the membrane of the nucleus-embedded vacuoles may have an altered identity. In support of this, when we tried to visualize the intra-nuclear vacuoles with MM4-64 and the nuclear periphery by using the ER marker Sec63-GFP in *yeh2Δ* cells treated with zeocin, we rarely detected any intranuclear vacuole. Only in isolated instances could we observe MM4-64 marks evocative of, yet inconclusively, intra-nuclear signals (Figure 3D).

To provide more solid proof for the identity of nuclear holes as vacuoles, we further used a dye marking the vacuolar lumen. To reduce the impact of the possibility that the lumen of intra-nuclear vacuoles also possesses altered properties, we chose the pH-independent dye BCECF (30). This robust dye allowed us to detect eventual examples where the nuclear hole was irrefutably filled with BCECF signals (Figure 3E, yellow arrowheads). Yet, in many instances, the BCECF signals arising from nuclear holes were poor or lacking, again suggesting that the engulfed vacuoles have altered properties. We conclude that the black holes detected in the nucleus most probably correspond to vacuoles whose identity is altered.

### 6. Alterations in autophagy are detected in cells with engulfed vacuoles

The vacuole is the central organelle where autophagy takes place. Vacuole sequestration in the nucleus could therefore alter the efficiency of autophagy. In particular, we hypothesized that their internalization in the nucleus would prevent them from interacting with the different cargoes, thus decreasing autophagy efficiency. To monitor this, we used a broadly accepted tool consisting of the N-terminally GFP-tagged version of the autophagosome membrane-nucleating factor Atg8 (31,32). GFP-Atg8 molecules become degraded with the cargo, but the partial resistance to degradation of the GFP moiety permits the assessment of autophagy completion. This way, degradative vacuoles appear as green when scored by fluorescence microscopy. Additionally, free GFP molecules, which migrate faster in a protein gel than the intact GFP-Atg8 ones, can be used to establish the percentage of degradation by Western blot. We compared the autophagic flux in conditions displaying increasing levels of nuclear holes (WT < *yeh2Δ* < WT + zeo < *yeh2Δ* + zeo, Figure 2G). We first treated or not WT and *yeh2Δ* cells with zeocin (zeocin effect was monitored by its ability to elicit the phosphorylation of the DNA Damage Response effector Rad53 (Figure S2C)), and induced autophagy 210 min later by adding rapamycin. We observed a striking correlation between nuclear holes presence and autophagic flux. Yet, to our surprise, it was inverse to our expectations: the conditions triggering the higher number of cells with nuclear holes were the ones showing increased autophagic completion, irrespective of whether monitoring was done by Western blot (Figure 4A, % free GFP moieties) or by counting green vacuoles (Figure 4B). Thus, scenarios in which nuclear holes are formed match a higher autophagic flux upon rapamycin induction.

**Figure 4.**
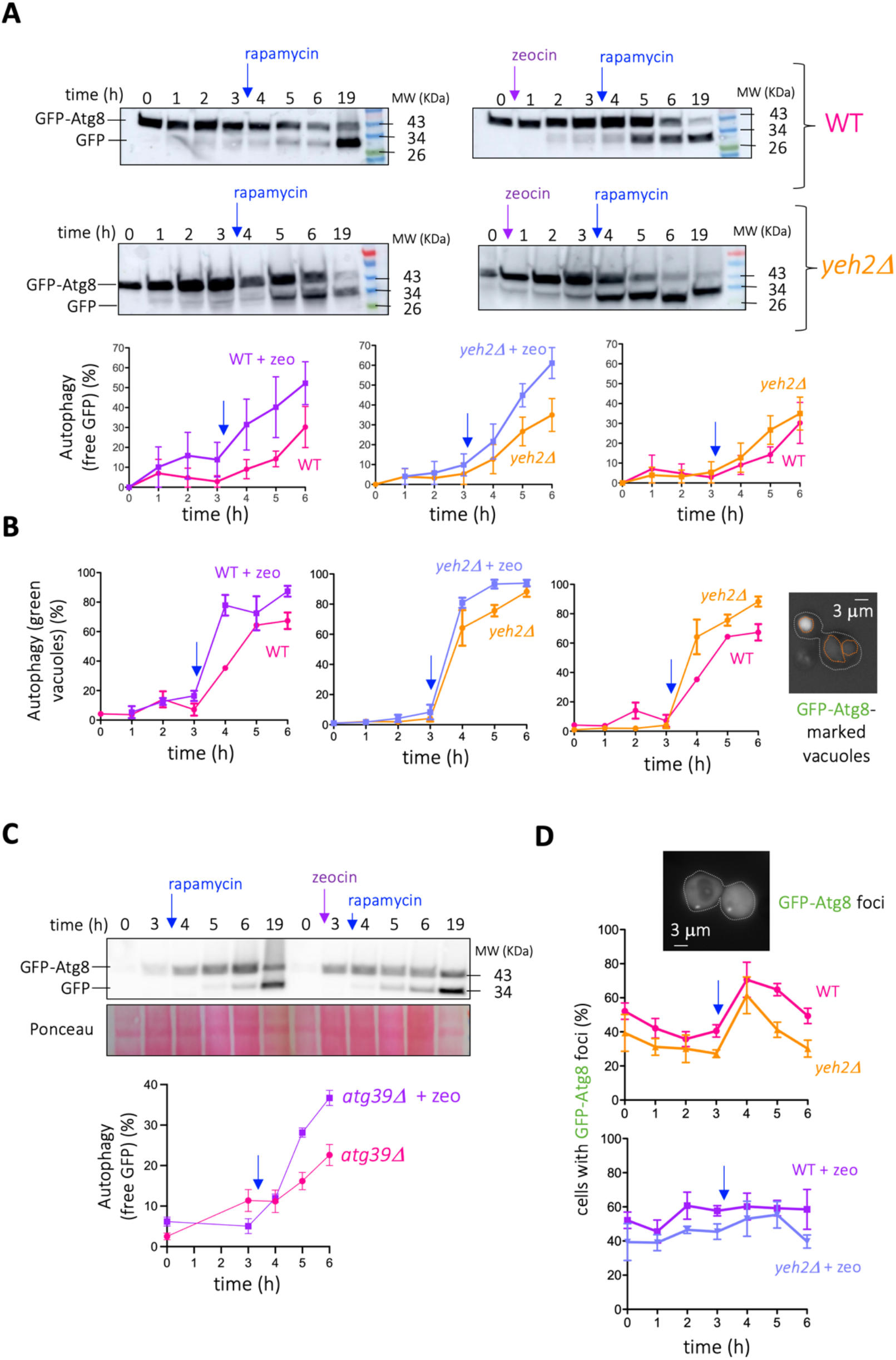
Characterization of the relationship between nucleus-internalized vacuoles and autophagy. **(A)** Cells of the indicated genotype, transformed with a vector expressing GFP-Atg8, were grown to the exponential phase in rich medium and treated as indicated. Notably, 100 μg/mL zeocin was added or not and, 3 h later, 200 ng/mL rapamycin was added in all the cases. Samples were retrieved at the indicated time-points. The implementation of autophagy was monitored through Western Blotting against GFP moieties. Time 16h is included to illustrate that cells achieve a comparable level of autophagy. The quantifications plot the percentage of free GFP with respect to all the GFP signal in a given lane. The blue arrow is a reminder of the moment when rapamycin was added. The plotted points and the error bars are the mean and the SEM, respectively, of 3 independent kinetics. **(B)** The same experiment described in (A) was done and cells were monitored by fluorescence microscopy. In this case, the level of autophagy was calculated as the percentage of cells in the population displaying green vacuoles, indicative of autophagy completion. An example of such a cell is displayed on the right, with the cell contour drawn in white and the vacuole one in orange. The blue arrow is a reminder of the moment when rapamycin was added. The plotted points and the error bars are the mean and the SEM, respectively, of 3 independent kinetics. **(C)** Details as in (A) but to compare the effect of zeocin treatment (or its absence) when autophagy is induced by rapamycin in a strain lacking the protein Atg39. **(D)** The same experiments presented in (B) were exploited to count the percentage of cells displaying GFP-Atg8 foci, indicative of growing phagophores and of autophagosomes. An example of a cell displaying two of these foci is shown, with the cell contour marked with a dashed white line.

### 7. Nucleus-internalized vacuoles represent low-efficiency vacuoles that basally tame autophagy if in the cytoplasm

We failed to detect nuclear membrane markers (e.g. Sec63-GFP signals) surrounding internalized vacuoles (Figure 3D). Yet, one could imagine that an increase in nuclear membrane may accompany the internalization. In that case, a trivial explanation underlying the observed increase in autophagy would be that the increase in nuclear membranes developed during vacuole engulfment provides more substrates to be degraded by autophagy. To assess this possibility, we repeated the monitoring of GFP-Atg8 degradation by comparing the ability in untreated *versus* zeocin conditions of a strain in which the Atg39 receptor, which instructs the autophagy of the nuclear membrane (33,34), had been removed. We observed that the incapacity to execute nuclear membrane autophagy still granted an accelerated autophagy completion when plus zeocin (Figure 4C). Thus, the enhanced autophagy execution seen in contexts where vacuoles are internalized in the nucleus does not stem from an excess of nuclear membrane-derived cargoes.

Still, the increase in general autophagy completion could be arguably explained by an increase in the number of cargoes to be degraded other than from the nuclear membrane. To assess this, we counted the number of cells displaying GFP-Atg8 foci, which represent the incipient phagophores and the growing and mature autophagosomes prior to their fusion to and destruction within the vacuole. This way, one would expect more foci in *yeh2Δ* than in WT cells if there were more cargoes to be degraded, while one would expect less foci in *yeh2Δ* than in WT cells if the increased autophagy completion was due to more efficient autophagosome clearance. Both when zeocin was added and when not, *yeh2Δ* cells recurrently displayed less GFP-Atg8 foci than WT cells (Figure 4D). Altogether, we conclude that the frequency with which the nucleus internalizes vacuoles matches an improvement in the ability of executing autophagy upon rapamycin addition. In this context, we propose that engulfed vacuoles may represent entities with low autophagy efficiency whose internalization within the nucleus would help cytoplasmic cargoes increase their chances of encountering only the proficient ones (Figure 5).

**Figure 5.**
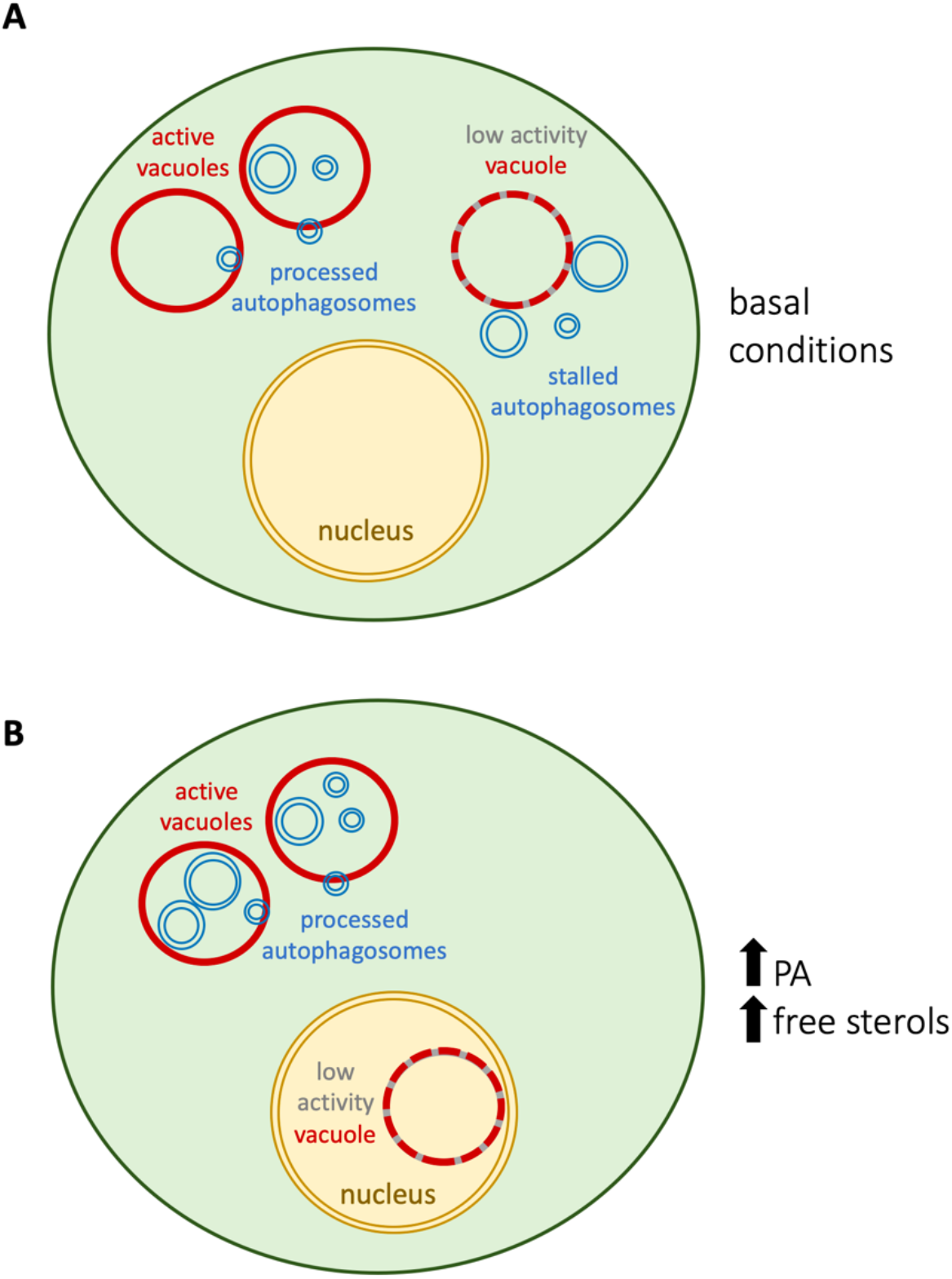
Proposed model to account for the improvement in autophagy in vacuole-engulfed contexts. **(A)** We propose that, basally, vacuoles with low autophagic potential may exist in the cytoplasm. Docking of autophagosome with such vacuoles does not culminate with autophagy execution yet delays these autophagosomes from delivering their content to a fully proficient vacuole. **(B)** When low autophagic potential vacuoles are internalized in the nucleus, for example in PA- or sterols-rich membrane scenarios, their clearance from the cytoplasm indirectly contributes to the more frequent encounter between proficient vacuoles and autophagosomes, therefore permitting an optimization of the autophagic flux.

## Discussion

In this work, we have identified a striking phenomenon through which the vacuole appears internalized in the nucleus. This process necessitates the accumulation of Phosphatidic Acid at the inner nuclear membrane and can be further fostered by high levels of free sterols. While its specific drivers and the detailed underlying process remain to be assessed in more depth, we uncover its impact on the efficiency of autophagy. Counterintuitively, sequestration of vacuoles in the nucleus matches an improved capacity for autophagy, presumably because low-efficiency vacuoles are the ones being internalized. Thus, we have uncovered a dramatic membrane-remodeling event with an immediate impact on metabolic adaptation.

We have serendipitously identified two situations in which, upon treatment of cells with two genotoxic agents, the phenomenon of vacuole internalization within the nucleus could be detected. It is hard to establish a common feature that can explain this, since zeocin and MMS, the two triggering agents, do not create DNA damage in the same way. In this sense, zeocin provokes single and double DNA breaks, while MMS mostly alkylates DNA bases and therefore limits the use of DNA as a template. Further, other genotoxins do not elicit the phenomenon under study (Figure 1B, 1SB). Thus, we think we can safely say that the phenomenon is not related to DNA damage itself. The explanation could be temptingly related to the cell cycle phase, since zeocin forces cells to arrest in G2 (Figure 1C) and nocodazole, an agent forcing cells to accumulate in the G2-to-M transition, also firmly elicits the phenotype (Figure 1D). That said, not all the treatments that lead cells to stall their progression at this cell cycle stage induce the formation of nuclear holes (Figure 1B,C). We purport that all the eliciting agents may entail a stress provoking changes in the metabolism of lipids. In support, zeocin was reported to trigger membrane expansion (23) and a recent work reports on methylglyoxal inducing a lipid-driven nuclear deformation after vacuolar pushing (35).

The mechanism through which vacuole ingression occurs also remains unassessed, but we define the need for a raise in the level of PA or sterols. Given the negative curvature-imparting potential of these lipids, in a first scenario the nuclear membrane would be plastically deformed towards the nucleoplasm to permit the vacuole entry. Resealing of the nuclear membranes behind the vacuole would sequester it without necessarily hosting it within the nucleoplasm. This option is unlikely because we failed to find intra-nuclear Sec63-GFP signals surrounding the vacuoles. Alternatively, the nuclear membrane may transiently break and reseal to permit a true engulfment. Yet, PA is also known as a fusogenic lipid (36), and the altered identity of the vacuole membrane once inside the nucleus (Figure 3D) is evocative of a vacuolar-nuclear membranes fusion event. Both sterols and PA have been defined as modulators of vacuolar membranes fusion (37,38), thus this scenario is also plausible. Whichever the case, the process will imply a deep remodeling of the contacts between the nuclear and vacuolar membranes. It is therefore possible that proteins implicated in vacuole-nucleus membranes hyper-tethering, such as Nvj1, Mdm1 and Snd3, are important players (5,8,9,15). Their study will be a key entry point to dissect the mechanism of vacuole ingression in the future.

The intrusion in the nucleoplasmic space of a voluminous body, such as a vacuole, is akin to alter nuclear processes. In a passive manner, just because of the space it occupies, it is likely to disrupt chromosome territories and displace, literally pushing, chromatin. Pushed chromatin behaves as condensed or compacted, and as such will emit related signals (39), impacting DNA transactions as transcription or repair. Furthermore, the presence of a membrane-enclosed body in the middle of the nucleus represents an unscheduled source of additional anchorage. Since multiple genome-related mechanisms, including coordination of transcription with replication, DNA damage sensing and repair, and telomere homeostasis request the nuclear membrane for structuration (40–46), the sudden presence of this novel substrate may interfere, for good or for bad, with these processes.

Our discovery of an increased ability for efficient autophagy under set-ups where some vacuoles were internalized in the nucleus raised the notion that vacuoles with low autophagic potential could be “selectively” excluded from the cytoplasm in this way (Figure 4). We suggest that this process may optimize the encounter of autophagosomes only with performant vacuoles, thus fostering the autophagic flux, quality control-wise (Figure 5, model). This model invokes a means for detecting the vacuoles whose autophagic potential is low. It is plausible that such vacuoles have an altered membrane composition that renders them susceptible of interaction with, and internalization within, the nucleus (Figure 5, discontinuous vacuole membrane). Our model also raises the question of whether this process could become “useful” and, as such, exploited by the cell under given circumstances. For example, the fact that the autophagy rate seen in the WT strain can be further improved by increasing sterols in membranes (Figure 4A,B, *yeh2Δ vs* WT) suggests that autophagy is basally “dampened” in the WT strain. Whether this window for autophagic capacity improvement is valuable under some circumstances remains to be explored. One of the set-ups where vacuole internalization within the nucleus was maximal corresponded to the absence of the phosphatase Nem1 (Figure 2C). In apparent contradiction with our proposal, absence of Nem1 hampers autophagy (47,48). We reconcile both findings claiming that, in the absence of Nem1, the internalization is so dramatic that it no further discriminates the type of vacuole, an extreme case in which the overall effect on autophagy will be negative.

Another aspect worth discussing is the potential conservation of this phenomenon. The vacuole fulfills in *S. cerevisiae* the function the lysosomes accomplish in most animal and vegetal cells. In these cells, a strikingly reminiscent process regarding the proximity of lysosomes with the nucleus dictates the cell’s ability to complete autophagy. Indeed, while autophagosomes form randomly at different locations within the cytoplasm, active lysosomes reside at the perinuclear region, and microtubule-dependent transport of autophagosomes towards the nuclear periphery needs to occur for fusion and subsequent autophagy (49). Of note, low cholesterol at membranes prevents this transport, therefore decreasing autophagy execution (50). Together, a common picture emerges in which, the higher the proximity with the nucleus (in the case of yeast being extreme, for it can be “inside”), the higher the overall efficiency in autophagy, and in both cases improved by high levels of free cholesterol in membranes. Indeed, lysosome positioning affects its acidity and therefore its autophagic potential (51–53), another aspect we also evoked during this work. In further (striking) analogy, a very recent work reports the accumulation of non-functional lysosomes within nuclear holes in late stages of the cell cycle in human cells (54). Last, the proximity of lysosomes to the nucleus is said to favor the faster delivery of transcription factors that reside onto lysosomes in order to trigger adaptive transcriptional programs in view of nutritional and metabolic changes (55). We note that hosting the vacuole literally inside the nucleus is the most radical way of bringing its coating transcription factors in proximity to their target DNA. It would be worth exploring whether we have uncovered, through this mechanism, a novel strategy for transcriptional regulation.

## Materials and Methods

### Cell culture and treatments

*Saccharomyces cerevisiae* cells were grown at 25°C in YEP (rich) or yeast nitrogen base (YNB) (minimal) liquid medium supplemented with 2% glucose (dextrose), unless otherwise indicated. Transformed cells were selected for plasmid maintenance in YNB– leucine or YNB–uracil medium overnight. The morning after, the exponentially growing cultures were diluted and grown for at least 4 hours in rich medium to create the optimal conditions to induce the formation of the “nuclear holes”, unless otherwise indicated. To induce the overexpression of *DGK1* and *PAH1-7A*, cells were grown overnight in YNB–leucine with 2% glycerol. Then, 2% galactose was added to exponentially growing cultures to induce their expression. The strains and the plasmids used in this study are referred to in Table 1 and Table 2, respectively.

**Table 1.**
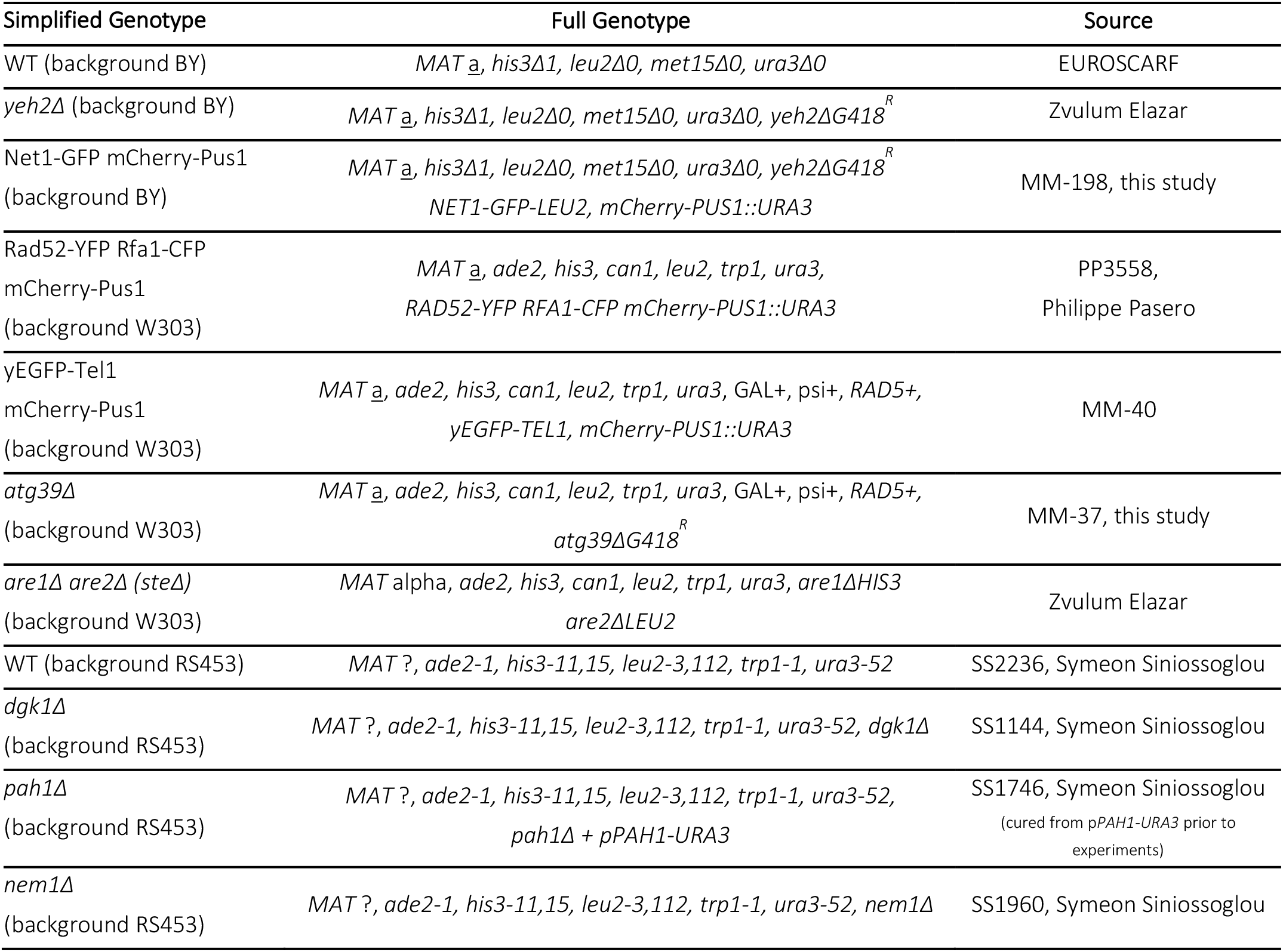
Strains used in this study

**Table 2.** Plasmids used in this study

| Simplified Name | Detailed Information | Source |
| --- | --- | --- |
| pEmpty (-ura) | pRS316 | Benjamin Pardo |
| pEmpty (-leu) | YEplac181 | Symeon Siniosoglou (16) |
| pDGK1 | YEplac181-GAL1/10p-DGK1 | Symeon Siniosoglou (17) |
| pPAH1-7A | YEplac181-GAL1/10p-PAH1-7A | Symeon Siniosoglou (16) |
| pSEC63-GFP | pJK59(CEN-URA3-SEC63p-SEC63-GFP(S65T,V163A)) | Sebastian Schuck (18) |
| pNOP1-CFP | pNOP1-CFP::LEU2 | Danesh Moazed (19) |
| pNET1-GFP | pDM266 (pNET1-GFP-LEU2) non-centromeric,<br>digested with *Bgl*II allows integration within the endogenous *NET1* locus | Félix Machín (20) |
| pNLS-Q2-mCherry | pRS316-CYC1p-Nup60<sup>1-24</sup>(NLS)-Opi1<sup>Q2</sup>-mCherry-NUP1t | Alwin Köhler (21) |
| pGFP-Atg8 | pGFP-ATG8-URA3 | Wei-Pang Huang (22) |

### Reagents

4-NQO (N8141, Sigma-Aldrich), rapamycin (HY-10219, Cliniscience), MM4-64 (SC-477259, Santa Cruz Biotechnology), BCECF (216254, Sigma-Aldrich), methylmetanosulfonate (MMS, 129925, Sigma-Aldrich), zeocin (R25001, ThermoFisher), Nocodazole (M1404, Sigma-Aldrich), Hydroxyurea (HU, H8627, Sigma-Aldrich), Camptothecin (CPT, C9911, Sigma-Aldrich).

### Cytometry

430 μL of culture samples at 10^7^ cells/mL were fixed with 1 mL of 100% ethanol. Cells were centrifuged for 1 minute at 16000g and resuspended in 500 μL 50 mM Na-Citrate buffer containing 5 μL of RNase A (10 mg/mL, Euromedex, RB0474) for 2 hours at 50°C. 6 μL of Proteinase K (Euromedex, EU0090-C) were added for 1 hour at 50°C. Aggregates of cells were dissociated by sonication (one 3 s-pulse at 50% potency in a Vibracell 72405 Sonicator). 20 μL of this cell suspension were incubated with 200 μL of 50 mM Na-Citrate buffer containing 4 μg/mL Propidium Iodide (FisherScientific). Data were acquired and analyzed on a Novocyte Express (Novocyte).

### Protein Extraction & Western blot

Approximately 5 × 10^8^ cells were collected at each relevant time point and washed with 20% trichloroacetic acid to prevent proteolysis, then resuspended in 200 μL of 20 % trichloroacetic acid at 4°C. The same volume of glass beads was added, and cells were disrupted by vortexing for 10 min. The resulting extract was spun for 10 min at 1000 g also at room temperature and the resulting pellet resuspended in 200 μL of 2x Laemmli buffer. Whenever the resulting extract was yellow-colored, the minimum necessary volume of 1 M Tris base (non-corrected pH) was added till blue color was restored. Then, water was added till reaching a final volume of 300 μL. These extracts were boiled for 10 min and clarified by centrifugation as before. To separate Rad53 isoforms, 10–15 μL of this supernatant was loaded onto a commercial 3–8% acrylamide gradient gel (BioRad) and migrated 70 min at 150 V in 1x Tris-Acetate buffer. The same volume of supernatant was used to separate GFP from GFP-Atg8 isoforms onto a commercial 4–20% acrylamide gradient gel (BioRad) and migrated 45 min at 100 V in 1x MES buffer. Proteins were transferred to a nitrocellulose membrane. Detection by immunoblotting was accomplished with anti-Rad53 antibody (1/3000), a kind gift from Dr. C. Santocanale, Galway, Ireland; or anti-GFP antibody (TP-401, Clinisciences, 1/2000), respectively, and in both cases an anti-rabbit HRP secondary antibody (A9044-2ML, Merck, 1/3000).

### Microscopy

1 mL of the culture of interest was centrifuged; then, the supernatant was thrown away and the pellet was resuspended in the remaining 50 μL. Next, 3 μL of this cell suspension was directly mounted on a coverslip for immediate imaging of the pertinent fluorophore-tagged protein signals. To dye vacuole membranes, 2 μL of a 4 mM MM4-64 stock were added to 1 mL of culture under incubation 30 min before visualization. To dye vacuole lumens, BCECF was added to and mixed with the the 50 μL of centrifuged pellet with residual medium at a 50 μM final concentration immediately prior to mounting. Imaging was achieved using a Zeiss Axioimager Z2 microscope and visualization, co-localization, and inspection performed with Image J.

### Quantification of Western blots

Image J was used to determine the pixel intensity values associated with the two bands (GFP-Atg8 and GFP) present in each lane. The percentage of autophagy was calculated by dividing the signal associated to free GFP divided the total signal measured in the lane, multiplied by 100.

### Quantification of Images

the determination of the percentage of cells in the population displaying nuclear holes was done by visual counting by the experimenter. Three independent experimenters participated in this counting as to warrant reproducibility and reliability.

### Graphical representations and Statistical analyses

were made with GraphPad Prism to both plot graphs and statistically analyze the data. For data representation, the SEM (standard error of the mean) was used. The SEM estimates how far the calculated mean is from the real mean of the sample population, while the SD (standard deviation) informs about the dispersion (variability) of the individual values constituting the population from which the mean was drawn. Since all the measurements we were considering for each individual experiment concerned a mean (the percentage of cells in the population presenting nuclear “holes”), and the goal of our error bars was to describe the uncertainty of the true population mean being represented by the sample mean, we did the choice of plotting the SEM.

## Acknowledgements

We are very thankful to Sebastian Schuck for the gift of pSec63-GFP, to Symeon Siniossoglou for the plasmid allowing mCherry-Pus1 tagging, the vectors to overexpress Dgk1 and Pah1-7A, and the *dgk1Δ, nem1Δ* and *pah1Δ* strains, and to Zvulum Elazar for *steΔ* and *yeh2Δ* strains. We also thank Alwin Köhler for pNLS-Q2-mCherry, Félix Machín for pNet1-GFP, Danesh Moazed for pNop1-CFP, Alba Torán-Vilarrubias for creating the *atg39Δ* strain, Corrado Santocanale for the kind gift of anti-Rad53 antibody, Pr. Wei-Pang Huang for the present of pGFP-Atg8, and Philippe Pasero and Benjamin Pardo for the Rad52-YFP Rfa1-CFP strain and pRS316, respectively. We are indebted to Simonetta Piatti, Lucile Espert and Félix Machín for critical reading of the manuscript and helpful insights to improve this work. We also acknowledge the imaging facility MRI, a member of the national infrastructure France-BioImaging, supported by the French National Research Agency (ANR-10-INBS-04, Investissements d’avenir). We finally thank the ATIP-Avenir program, La Ligue contre le Cancer et l’Institut National du Cancer (PLBIO19-098 INCA_13832), France, for funding our research.

## Author contributions

Conceptualization, M.G., S.K., A. E.-V. and M.M.-C.; Data curation, S.K., A. E.-V. and M.M.-C.; Formal analysis, M.G., S.K., A. E.-V., C. S. and M.M.-C.; Methodology, M.G., S.K. and M.M.-C.; Investigation, M.G., S.K. and M.M.-C.; Writing—original Draft, M.M.-C.; Writing—Review and Editing, M.G., S.K., A. E.-V., C.S. and M.M.-C.; Funding Acquisition, M.M.-C.; Supervision, M.M.-C. Project administration, M.M.-C.

## Abbreviations

BCECF: 2’,7’-Bis-(2-Carboxyethyl)-5-(and-6)-Carboxyfluorescein Acetoxymethyl Ester
CFP: cyan fluorescent protein
CPT: camptothecin
DAG: diacylglycerol
DIC: differential interference contrast
GFP: green fluorescent protein
HU: hydroxyurea
INM: inner nuclear membrane
MMS: methylmethanosulfonate
NVJ: nucleus-vacuole junction
PA: phosphatidic acid
SEM: standard error of the mean
WT: wild type
4-NQO: 4-nitroquinoline 1-oxide.

---

**Figure S1.**
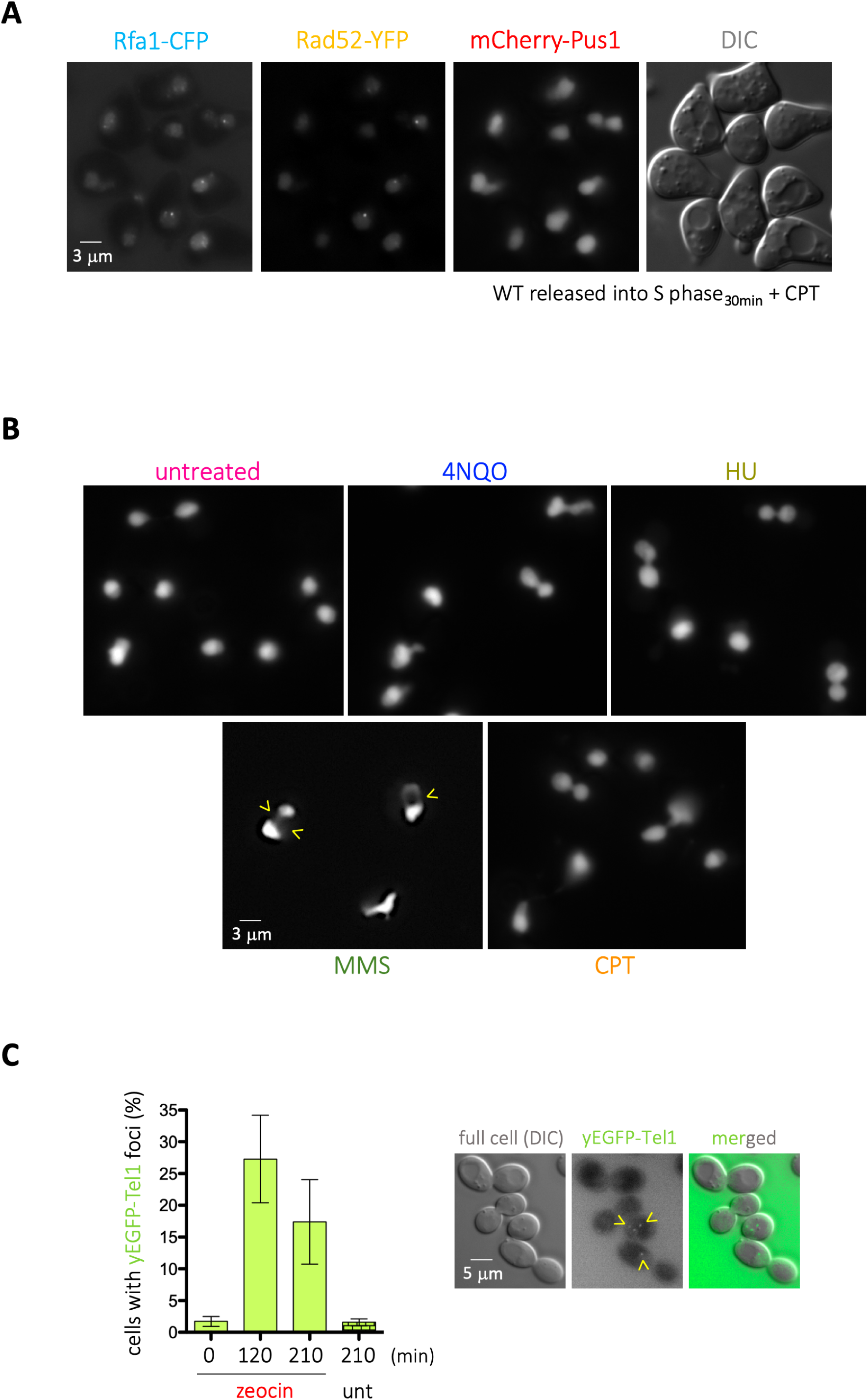
Additional information related to Figure 1. **(A)** Illustrative images of a canonical experiment in which we monitor the ability of DNA repair proteins (Rad52-YFP, Rfa1-CFP) to form foci using mCherry-Pus1 as a marker to define the nucleus boundaries. **(B)** Illustrative images of nuclei, as revealed by the mCherry-Pus1 signals, in response to the different genotoxins used in Figure 1. Deformations reminiscent to holes are indicated by yellow arrowheads. **(C)** Quantification of the percentage of cells displaying yEGFP-Tel1 foci in response to 100 μg/mL zeocin while growing in minimal medium at the indicated times. One illustrative image of the events being counted is shown.

**Figure S2.**
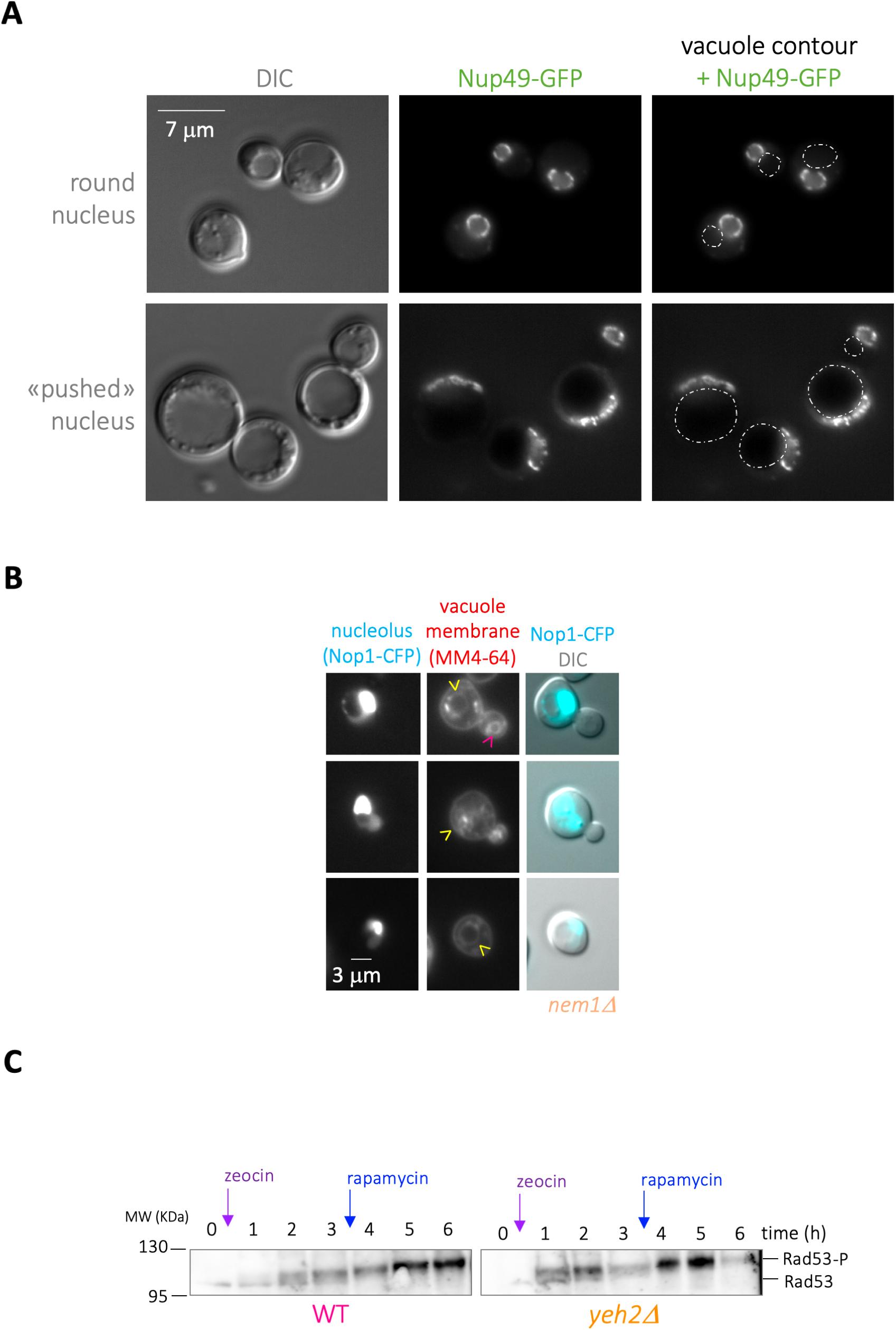
Additional information related to Figures 3 & 4. **(A)** Cells bearing a GFP-tagged Nup49 in order to define the nuclear periphery were pictured at two moments during growth, namely at late exponential phase (top) or after nutrient exhaustion (bottom). The vacuole contour, as drawn from the DIC images, is over-imposed on the Nup49-GFP images to appreciate how it can, when enlarged, push nuclei. **(B)** The *nem1Δ* strain, transformed with the vector expressing Nop1-CFP, was simultaneously dyed with the vacuole membrane marker MM4-64. Nop1-CFP signals are overexposed to allow the visualization of the nuclear hole, mostly present in the non-Nop1-marked part of the nucleus. The MM4-64 signal coming from the hole-residing vacuole is poor (yellow arrowheads), and contrasts with that of the MM4-64 signal coming from cytoplasmic vacuoles (pink arrowhead). **(C)** The same zeocin-treated samples used for Western Blot in Figure 4A were used here to monitor (and validate) the activation of the DNA Damage Response effector Rad53, which can be visualized as a progressive phosphorylation since the addition of zeocin.

## References

1. Ron D, Walter P. Signal integration in the endoplasmic reticulum unfolded protein response. Nat Rev Mol Cell Biol. 2007;8(7):519–29.

2. Zhang D, Oliferenko S. Remodeling the nuclear membrane during closed mitosis. Curr Opin Cell Biol [Internet]. 2013;25(1):142–8. Available from: http://dx.doi.org/10.1016/j.ceb.2012.09.001

3. Ungricht R, Kutay U. Mechanisms and functions of nuclear envelope remodelling. Nat Rev Mol Cell Biol. 2017;18(4):229–45.

4. Scorrano L, De Matteis MA, Emr S, Giordano F, Hajnóczky G, Kornmann B, et al. Coming together to define membrane contact sites. Nat Commun [Internet]. 2019;10(1):1–11. Available from: http://dx.doi.org/10.1038/s41467-019-09253-3

5. Rogers SM, Hariri H, Wood NEM, Speer NO, Henne WM. Glucose restriction drives spatial reorganization of mevalonate metabolism. Elife. 2021;10:1–23.

6. Lahiri S, Toulmay A, Prinz WA. Membrane contact sites, gateways for lipid homeostasis. Curr Opin Cell Biol [Internet]. 2015;33:82–7. Available from: http://dx.doi.org/10.1016/j.ceb.2014.12.004

7. Pan X, Roberts P, Chen Y, Kvam E, Shulga N, Huang K, et al. Nucleus-vacuole junctions in Saccharomyces cerevisiae are formed through the direct interaction of Vac8p with Nvj1p. Mol Biol Cell. 2000;

8. Tosal-Castano S, Peselj C, Kohler V, Habernig L, Berglund LL, Ebrahimi M, et al. Snd3 controls nucleus-vacuole junctions in response to glucose signaling. Cell Rep. 2021;34(3).

9. Hariri H, Rogers S, Ugrankar R, Liu YL, Feathers JR, Henne WM. Lipid droplet biogenesis is spatially coordinated at ER –vacuole contacts under nutritional stress. EMBO Rep. 2018;19(1):57–72.

10. Kvam E, Goldfarb DS. Nucleus-vacuole junctions and piecemeal microautophagy of the nucleus in S. cerevisiae. Autophagy. 2007;3(2):85–92.

11. Mostofa MG, Morshed S, Shibata R, Takeichi Y, Rahman MA, Hosoyamada S, et al. rDNA Condensation Promotes rDNA Separation from Nucleolar Proteins Degraded for Nucleophagy after TORC1 Inactivation. Cell Rep [Internet]. 2019;28(13):3423–3434.e2. Available from: https://doi.org/10.1016/j.celrep.2019.08.059

12. Golam Mostofa M, Rahman MA, Koike N, Yeasmin AMST, Islam N, Waliullah TM, et al. CLIP and cohibin separate rDNA from nucleolar proteins destined for degradation by nucleophagy. J Cell Biol. 2018;217(8):2675–90.

13. Lord CL, Wente SR. Nuclear envelope-vacuole contacts mitigate nuclear pore complex assembly stress. J Cell Biol. 2020;

14. Kvam E, Goldfarb DS. Nucleus-vacuole junctions in yeast: Anatomy of a membrane contact site. Biochem Soc Trans. 2006;34(3):340–2.

15. Henne WM, Zhu L, Balogi Z, Stefan C, Pleiss JA, Emr SD. Mdm1/Snx13 is a novel ER-endolysosomal interorganelle tethering protein. J Cell Biol. 2015;210(4):541–51.

16. O’Hara L, Han GS, Sew PC, Grimsey N, Carman GM, Siniossoglou S. Control of phospholipid synthesis by phosphorylation of the yeast lipin Pah1p/Smp2p Mg2+-dependent phosphatidate phosphatase. J Biol Chem. 2006;281(45):34537–48.

17. Karanasios E, Barbosa AD, Sembongi H, Mari M, Han GS, Reggiori F, et al. Regulation of lipid droplet and membrane biogenesis by the acidic tail of the phosphatidate phosphatase Pah1p. Mol Biol Cell. 2013;

18. Prinz WA, Grzyb L, Veenhuis M, Kahana JA, Silver PA, Rapoport TA. Mutants affecting the structure of the cortical endoplasmic reticulum in Saccharomyces cerevisiae. J Cell Biol. 2000;150(3):461–74.

19. Mekhail K, Seebacher J, Gygi SP, Moazed D. Role for perinuclear chromosome tethering in maintenance of genome stability. Nature. 2008;

20. Matos-Perdomo E, Machín F. The ribosomal DNA metaphase loop of Saccharomyces cerevisiae gets condensed upon heat stress in a Cdc14-independent TORC1-dependent manner. Cell Cycle. 2018;17(2):200–15.

21. Romanauska A, Köhler A. The Inner Nuclear Membrane Is a Metabolically Active Territory that Generates Nuclear Lipid Droplets. Cell. 2018;

22. Wang SH, Lin PY, Chiu YC, Huang JS, Kuo YT, Wu JC, et al. Curcumin-mediated HDAC inhibition suppresses the DNA damage response and contributes to increased DNA damage sensitivity. PLoS One. 2015;10(7):1–19.

23. Witkin KL, Chong Y, Shao S, Webster MT, Lahiri S, Walters AD, et al. The budding yeast nuclear envelope adjacent to the nucleolus serves as a membrane sink during mitotic delay. Curr Biol. 2012;

24. Choudhary V, Golani G, Joshi AS, Cottier S, Schneiter R, Prinz WA, et al. Architecture of Lipid Droplets in Endoplasmic Reticulum Is Determined by Phospholipid Intrinsic Curvature. Curr Biol. 2018;

25. Ben M’barek K, Ajjaji D, Chorlay A, Vanni S, Forêt L, Thiam AR. ER Membrane Phospholipids and Surface Tension Control Cellular Lipid Droplet Formation. Dev Cell. 2017;

26. Müllner H, Deutsch G, Leitner E, Ingolic E, Daum G. YEH2/YLR020c encodes a novel steryl ester hydrolase of the yeast Saccharomyces cerevisiae. J Biol Chem. 2005;

27. Campbell JL, Lorenz A, Witkin KL, Hays T, Loidl J, Cohen-Fix O. Yeast nuclear envelope subdomains with distinct abilities to resist membrane expansion. Mol Biol Cell. 2006;

28. Walters AD, Amoateng K, Wang R, Chen JH, McDermott G, Larabell CA, et al. Nuclear envelope expansion in budding yeast is independent of cell growth and does not determine nuclear volume. Mol Biol Cell. 2019;30(1):131–45.

29. Barbosa AD, Lim K, Mari M, Edgar JR, Gal L, Sterk P, et al. Compartmentalized Synthesis of Triacylglycerol at the Inner Nuclear Membrane Regulates Nuclear Organization. Dev Cell [Internet]. 2019;50(6):755–766.e6. Available from: https://doi.org/10.1016/j.devcel.2019.07.009

30. Plant PJ, Manolson MF, Grinstein S, Demaurex N. Alternative mechanisms of vacuolar acidification in H+-ATPase-deficient yeast. J Biol Chem. 1999;274(52):37270–9.

31. Nair U, Thumm M, Klionsky DJ, Krick R. GFP-Atg8 protease protection as a tool to monitor autophagosome biogenesis. Autophagy. 2011;7(12):1546–50.

32. Geng J, Baba M, Nair U, Klionsky DJ. Quantitative analysis of autophagy-related protein stoichiometry by fluorescence microscopy. J Cell Biol. 2008;182(1):129–40.

33. Mochida K, Oikawa Y, Kimura Y, Kirisako H, Hirano H, Ohsumi Y, et al. Receptor-mediated selective autophagy degrades the endoplasmic reticulum and the nucleus. Nature. 2015;522(7556):359–62.

34. Otto FB, Thumm M. Mechanistic dissection of macro- and micronucleophagy. Autophagy. 2021;17(3):626–39.

35. Nomura W, Aoki M, Inoue Y. Methylglyoxal inhibits nuclear division through alterations in vacuolar morphology and accumulation of Atg18 on the vacuolar membrane in Saccharomyces cerevisiae. Sci Rep [Internet]. 2020;10(1):1–13. Available from: https://doi.org/10.1038/s41598-020-70802-8

36. Zhukovsky MA, Filograna A, Luini A, Corda D, Valente C. Phosphatidic acid in membrane rearrangements. FEBS Lett. 2019;593(17):2428–51.

37. Kato M, Wickner W. Ergosterol is required for the Sec18/ATP-dependent priming step of homotypic vacuole fusion. EMBO J. 2001;20(15):4035–40.

38. Miner GE, Starr ML, Hurst LR, Fratti RA. Deleting the DAG kinase Dgk1 augments yeast vacuole fusion through increased Ypt7 activity and altered membrane fluidity. Traffic. 2017;18(5):315–29.

39. Burgess RC, Burman B, Kruhlak MJ, Misteli T. Activation of DNA Damage Response Signaling by Condensed Chromatin. Cell Rep. 2014;

40. Bermejo R, Capra T, Jossen R, Colosio A, Frattini C, Carotenuto W, et al. The replication checkpoint protects fork stability by releasing transcribed genes from nuclear pores. Cell. 2011;

41. Towbin BD, Meister P, Gasser SM. The nuclear envelope - a scaffold for silencing? Current Opinion in Genetics and Development. 2009.

42. Rothballer A, Kutay U. The diverse functional LINCs of the nuclear envelope to the cytoskeleton and chromatin. Chromosoma. 2013.

43. Mekhail K, Moazed D. The nuclear envelope in genome organization, expression and stability. Nature Reviews Molecular Cell Biology. 2010.

44. Lei K, Zhu X, Xu R, Shao C, Xu T, Zhuang Y, et al. Inner nuclear envelope proteins SUN1 and SUN2 play a prominent role in the DNA damage response. Curr Biol. 2012;

45. Oza P, Jaspersen SL, Miele A, Dekker J, Peterson CL. Mechanisms that regulate localization of a DNA double-strand break to the nuclear periphery. Genes Dev. 2009;

46. Shibuya H, Hernández-Hernández A, Morimoto A, Negishi L, Höög C, Watanabe Y. MAJIN Links Telomeric DNA to the Nuclear Membrane by Exchanging Telomere Cap. Cell. 2015;163(5):1252–66.

47. Rahman MA, Terasawa M, Mostofa MG, Ushimaru T. The TORC1–Nem1/Spo7–Pah1/lipin axis regulates microautophagy induction in budding yeast. Biochem Biophys Res Commun. 2018;504(2):505–12.

48. Rahman MA, Mostofa MG, Ushimaru T. The Nem1/Spo7–Pah1/lipin axis is required for autophagy induction after TORC1 inactivation. FEBS J. 2018;285(10):1840–60.

49. Kimura S, Noda T, Yoshimori T. Dynein-dependent movement of autophagosomes mediates efficient encounters with lysosomes. Cell Struct Funct. 2008;33(1):109–22.

50. Wijdeven RH, Janssen H, Nahidiazar L, Janssen L, Jalink K, Berlin I, et al. Cholesterol and ORP1L-mediated ER contact sites control autophagosome transport and fusion with the endocytic pathway. Nat Commun. 2016;7(May).

51. Johnson DE, Ostrowski P, Jaumouillé V, Grinstein S. The position of lysosomes within the cell determines their luminal pH. J Cell Biol. 2016;212(6):677–92.

52. Gowrishankar S, Ferguson SM. Lysosomes relax in the cellular suburbs. J Cell Biol. 2016;212(6):617–9.

53. Korolchuk VI, Saiki S, Lichtenberg M, Siddiqi FH, Roberts EA, Imarisio S, et al. Lysosomal positioning coordinates cellular nutrient responses. Nat Cell Biol. 2011;13(4):453–62.

54. Almacellas E, Pelletier J, Day C, Ambrosio S, Tauler A, Mauvezin C. Lysosomal degradation ensures accurate chromosomal segregation to prevent chromosomal instability. Autophagy [Internet]. 2021;17(3):796–813. Available from: https://doi.org/10.1080/15548627.2020.1764727

55. Zhao Q, Gao SM, Wang MC. Molecular Mechanisms of Lysosome and Nucleus Communication. Trends Biochem Sci [Internet]. 2020;45(11):978–91. Available from: https://doi.org/10.1016/j.tibs.2020.06.004

56. Loewen CJR, Gazpar ML, Jesch SA, Delon C, Ktistakis NT, Henry SA, et al. Phospholipid metabolism regulated by a transcription factor sensing phosphatidic acid. Science (80) 2004

